# Spectral preferences of mosquitos are altered by odors

**DOI:** 10.1101/2025.02.05.636723

**Authors:** Adam J. Blake, Jeffrey A. Riffell

**Affiliations:** Department of Biology, University of Washington, Box 351800, Seattle, WA 98195-1800

**Keywords:** Insect vision, Mosquito, Olfaction, Multimodal Cues, Sensory integration, LED synth, Wind tunnel

## Abstract

Vision underlies many important behaviors in insects generally and in mosquitos specifically. Mosquito vision plays a role in predator avoidance, mate finding, oviposition, locating vertebrate hosts, and vectoring disease. Recent work has shown that when sensitized to CO2, the visual responses of *Aedes aegypti* are wavelength-dependent, but little is known about how other olfactory stimuli can modulate visual responses. The visual cues associated with flowers, vertebrate hosts, or oviposition sites differs substantially and it is possible that odors might prime the mosquito visual system to respond to these different resources. To investigate the interplay of olfactory and visual cues, we adapted previously used wind tunnel bioassays to use quasi-monochromatic targets (390-740 nm) created with a novel LED synth. We coupled these visual targets with CO2 and the odors representative of vertebrate hosts, floral nectar or oviposition sites and assessed responses via 3D tracking of female mosquitos. When CO2 alone is present, we observe a lower preference for wavelengths in the green portion of the visible spectrum with a gradual increase as wavelengths moved towards the violet and red ends of the spectrum. However, when odors associated both with flowers and oviposition sites, we observed significant increases in mosquito preference for green (475-575 nm) stimuli. In contrast when vertebrate host odor was present, we saw increased preference for stimuli across the entire visible spectrum. These odor shifts in the mosquito spectral preferences suggest these preferences are not fixed and shift depending on behavioral context.

## Introduction

Vision as a sensory modality plays an important role in many if not most ecological interactions among insects (Warrant 2019). Mosquitos, like most insects, use vision in a variety of ecological interactions, including predator avoidance, mate finding, oviposition, locating vertebrates hosts, and vectoring the pathogens of disease (Clements 1999; Hawkes et al. 2022). However, vision is used in concert with many other sensory cues including odor, heat, or humidity depending on the behavioral context. For example, in the context of vertebrate host finding in mosquitos, it has been shown that CO2 can induce visual search behaviors (van Breugel et al. 2015; Carnaghi et al. 2021). In the absence of CO2 mosquitos are not attracted towards dark high contrast objects, however after exposure to CO2 mosquitos will approach and investigate these objects. Other cues such as heat, odor, and humidity govern landing and biting (McMeniman et al. 2014; Cardé 2015; Sumner and Cardé 2022; Sumner et al. 2023; Giraldo et al. 2023). This CO2-gated approach of mosquitos to visual stimuli is wavelength dependent, with mosquitos attracted to the cyan, orange, and red spectral bands and showed some ability to discriminate between green and red stimuli of matched perceptual brightness (Alonso San Alberto et al. 2022).

Mosquitos have a visual system with many similarities with other dipterans (Hawkes et al. 2022), but we lack detailed information about their photoreceptors, unlike other flies such as *Calliphora*, *Musca,* or *Drosophilia* (Hardie 1985; Sharkey et al. 2020). Like these flies, mosquitos have compound eyes made up of hundreds of individual units know as ommatidium consisting of 6 outer (R1-6) and 2 inner (R7,8) photoreceptor cells (Brammer 1970). However, in higher flies opsin expression varies among ommatidial types in a random mosaic across most of the compound eye, whereas in mosquitos ommatidial types are highly regional with a single type in each eye region (Hu et al. 2009). Of the 10 opsins that have been identified in *Aedes aegypti*, 5 are expressed in the compound eye (Giraldo-Calderón et al. 2017). These include a pair of longwave green sensitive opsins, a blue and a UV sensitive opsin as well as opsin homologous with the Drosophila Rh7 that in mosquitos is also sensitive to green wavelengths (Hu et al. 2009; Hu et al. 2011; Hu et al. 2012; Hu et al. 2014). The outer photoreceptors always express the same green sensitive opsin while the expression of the central photoreceptors varies across the eye regions with the majority of the eye expressing green and UV sensitive opsins, with blue sensitivity limited to a small ventral stripe and the very dorsal portion of the eye. This opsin expression mirrors electroretinogram studies in *Ae. aegypti* showing two sensitivity peaks in the green (∼525 nm) and in the UV (∼350 nm) (Muir et al. 1992b). We lack sensitivity data for individual mosquito photoreceptors, and it is currently unclear if the multiple longwave sensitive opsins could provide the underpinnings to allow for the behavioral discrimination between green and red spectral bands.

Different odors have been shown to shift spectral preferences in other insects (Reisenman et al. 2000; Yoshida et al. 2015; Brodie et al. 2015; Balamurali et al. 2019; Bolton et al. 2021), which could be one reason for the conflicting results in behavioral studies examining spectral preference in mosquitoes. Many studies have identified red and black are attractive colors (Fay and Prince 1968; Muir et al. 1992a), including the recent study examining CO2 gating of visual responses (Alonso San Alberto et al. 2022), but other colors have also been found to be attractive (Brett 1938; Smart and Brown 1956; Snow 1971). If odors beyond CO2 could gate or shift the spectral preferences of mosquitos this would seem adaptive as reflectance spectra of resources important to mosquitos can differ considerable. For example, human skin reflects considerable amount of light in the 600-700 nm range (Stamatas et al. 2004) whereas the reflectance of many mosquito pollinated flowers is highest between 500-600 nm (Alonso San Alberto et al. 2022). It was then the objective of this study to investigate how other odors in addition to CO2 could alter the visual behavior of *Ae. aegypti*. To perform this objective, we adapted proved 3D tracking methods to characterize the movement of female mosquitos responding to visual stimuli in the presence of odors characteristic of human hosts, floral nectar sources, and potential oviposition sites. We refined earlier techniques to make use of novel LED synths (Belušič et al. 2016; Egri et al. 2020), allowing for much tighter control of stimulus wavelength and intensity, allowing for refined estimates of spectral preference independent of the confounding effects of stimulus intensity.

## Methods

### Photography and Spectroscopy

Photographic measurements (Blake and Riffell 2025) were taken with a Nikon D5100 DSLR (spectral sensitivity characterized in Jiang et al. 2013) fitted with a AF-S DX Nikkor 35mm f/1.8G lens. The captured raw images were decoded in FIJI (Schindelin et al. 2012), using the DCRAW plugin (Coffin 2019) in a manner that preserved sensor linearity.

Spectra (Figs. 1B, S1, Blake and Riffell 2025) were measured using a calibrated spectrophotometer (USB-2000, Ocean Optics Inc., Dunedin, FL, USA) with reflectance spectra being calibrated against a 99% Spectralon reflectance standard (SRS-99–010, Labsphere, NH).

**Figure 1.**
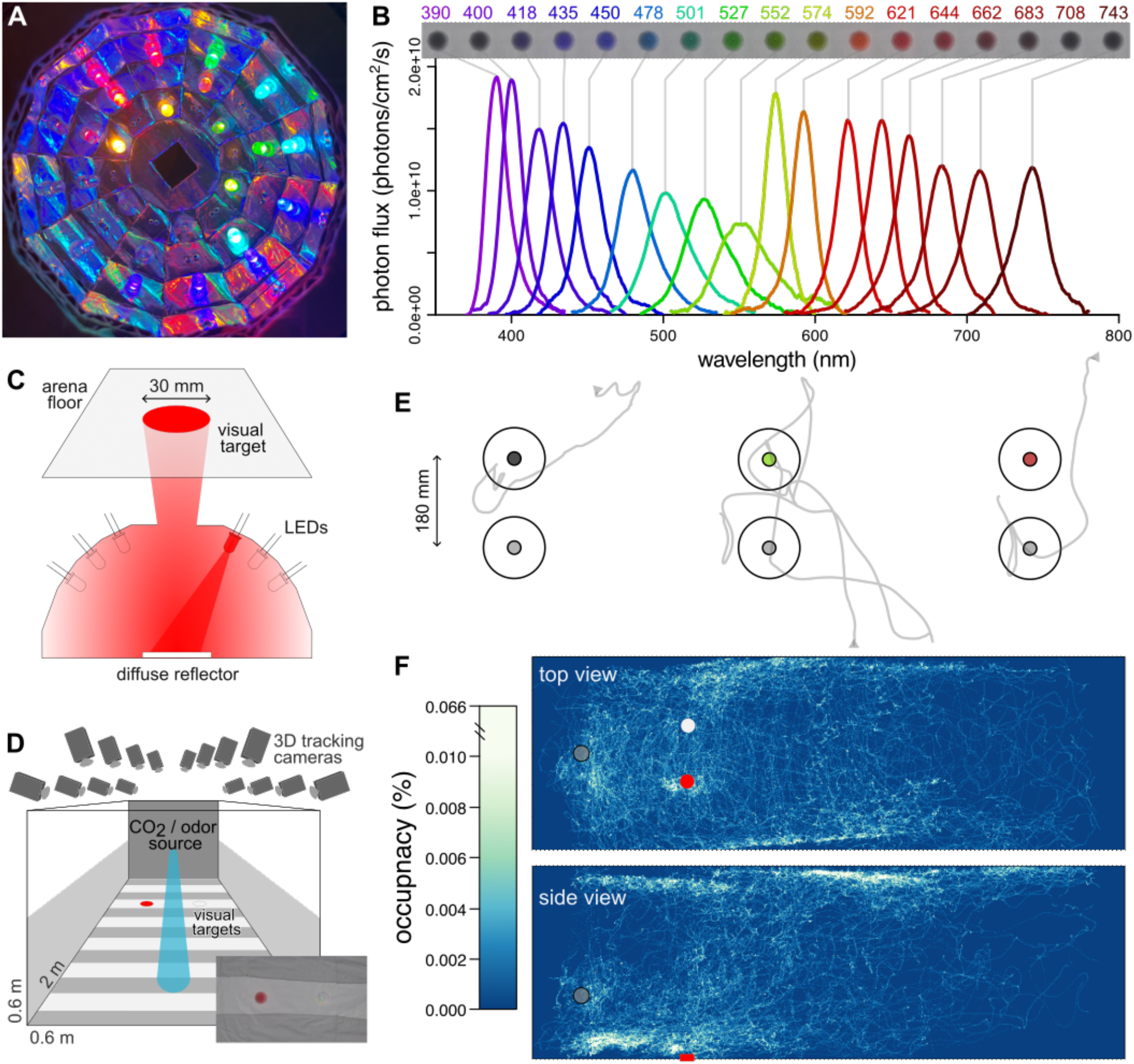
LED synth stimuli and wind tunnel bioassay. **A** Interior view of one of the LED synths showing the arrangement of LEDs. **B** Photon flux of the 17 LED color channels of the LED synths at isoquantal intensity used in the spectral sweep bioassays. **C** Diagram of an LED synth demonstrating how visual stimuli are generated through diffuse reflection. **D** Wind tunnel system showing position of visual stimuli, odor source, and tracking cameras. **(inset)** Photograph showing a top down view of visual stimuli. **E** Top down view of individual trajectories examples. The arrows represent the start of a trajectory; larger circles show the response cylinders, smaller circles are the visual stimuli. **F** Top and side view occupancy maps showing mosquito response to visual stimuli (red and white circles) and odor source (gray circle). The row of images above shows the appearance of these visual stimuli as they appear under conventional photography. The stimuli on either end of the visual spectrum appear dark as they fall outside of the camera’s (and human) spectral sensitivity.

### Experimental animals

Mosquitos (*Aedes aegypti*: Rockefeller) were provided from BEI Resources (Manassas, VA, USA) and were raised at the University of Washington campus in mixed sex groups of approximately 200 adults. Mosquitos were provided with sucrose *ad libitum* and maintained at 26–28 °C, 80% RH, and a photoperiod of 12L:12D. Females were used in behavioral experiments 7-8 days post emergence and were assumed to be mated. Previous research had established a mating rate of over 90% with cohabiting females of this age (Alonso San Alberto et al. 2022). Before use in experiments mosquitos were cold anesthetized, males removed, separated into groups of 35-50 females, and deprived of sucrose for a period of 16-24 hours.

Mosquitos were used in experiments during an approximately 6 h period centered around the mosquito’s subjective sunset. This time of day was chosen as mosquito flight activity increases in the approach of sunset and absent a lights off cue, this activity remains high for several hours following sunset (Clements 1999).

### Wind tunnel

These experiments made use of the same wind tunnel and real-time tracking system as described in previous papers (van Breugel et al. 2015; Vinauger et al. 2019; Zhan et al. 2021; Alonso San Alberto et al. 2022) and its construction and use are detailed in Alonso San Alberto et al. (2023). Any deviations from this setup and procedure are fully detailed below. Behavioral experiments took place in a low-speed wind tunnel (ELD Inc., Lake City, MN), with a working section of 224 by 61 by 61 cm high and a constant laminar flow of 40 cm/sec (Fig. 1D). We used two short-throw projectors (LG PH450U, Englewood Cliffs, NJ) and rear projection screens (SpyeDark, Spye, LLC, Minneapolis, MN; Fig S1A,C) to provide low contrast gray horizons on each side of the tunnel. The projectors provided ambient light at a level of ∼3 lux across the 420–670 nm range. Fabric liners were positioned on the floor of the working section to provide both a low contrast background and a projection surface for the LED synth arrays. The custom sewn liner consisted of a piece of white cotton broadcloth (Fig S1D,E, Jo-Ann Stores, LLC, Hudson, OH) with 10 strips of black tulle (Fig S1D,E, Jo-Ann Stores, LLC) appliquéd at regular intervals.

Sixteen cameras (Basler AC640gm, Exton, PA) fitted with IR pass filters (Kodak 89B, Kodak, Rochester, NY) were positioned so that multiple cameras covered all areas of the working section allowing mosquito trajectories to be recorded at 60 frames/sec. IR backlights (B07ZZ2LJKY, 360DigitalSignage, Shenzhen, GD, China) were installed below and the sides of the wind tunnel and diffused by the side screens and fabric floor to provide a uniform bright background in the IR to optimize mosquito tracking while falling well outside the visual sensitivity range of the mosquitoes (Fig S1B). The temperature within the working section was a constant 22.5°C and ambient CO2 outside the wind tunnel was ∼400 ppm (Alonso San Alberto et al. 2022).

### CO2 and odor stimuli

The CO2, filtered clean air, and odor-laden air were delivered using three mass flow controllers (MC-200SCCM-D, Alicat Scientific, Tucson, AZ) whose output were combined and delivered to the working section via a single outlet located upwind of the visual stimuli at a height of 20 cm (Fig. 1D). The mass flow controllers were supplied with clean filtered air or CO2 via gas canisters (Linde Gas & Equipment Inc., Burr Ridge, IL). The timing and flow rates of CO2, filtered clean air, and odor-laden air were independently controlled by the same Python script that controlled visual stimuli. All stimulus series included a pre-CO2 and post-CO2 period to serve as a baseline of mosquito behavior in the absence of attractive stimuli. Preliminary testing indicated that the CO2 concentration (0%, 1%, 5%, or 10%) in the plume had no effect on spectral preference but did significantly increase both activation and recruitment to visual stimuli (Fig S2). For these reasons, we elected to perform all bioassays using a plume with 10% CO2, despite this concentration being greater than what would be expected in the vicinity of vertebrate hosts (Geier et al. 1999), sources of floral nectar (Peach et al. 2019a), or oviposition sites.

Odor-laden air was directed through a 1 L container enclosing the odor source, with the odor, when present, composing 10% of air being released from the outlet. Odor sources were changed in between runs, and the odor containers were switched out and cleaned with 95% ethanol. Runs testing a floral odor used a freshly cut common tansy (*Tanacetum vulgare*) inflorescence, with 10–15 composite flowers and its stem inserted into a water-filled vial (20 ml). These flowers were collected from the vicinity of the University of Washington campus in Seattle, WA. Common tansy was chosen as its scent had been previously demonstrated to be attractive to *A. aegypti* (Peach et al. 2019b; Peach et al. 2019a).

Human foot odor has been shown to be behaviorally attractive to *A. aegypti* (Lacey et al. 2014; Bello and Cardé 2022) and is easily collected using nylon socks (Njiru et al. 2006; Okumu et al. 2010). In runs testing a human order, a single nylon sock was used as an odor source. The socks were worn for a period of 8-12 hours by one of the investigators (AJB). Odor collection was limited to a single individual to eliminate the potential of interindividual variability in odor composition (Ellwanger et al. 2021).

Plant infusions have been demonstrated to be effective oviposition attractants for *A. aegypti* (Reiter 1991; Mwingira et al. 2020). Following the methods of (Ritchie et al. 2014) we created an extract using 2 g of dried alfalfa in 1.2 L of water that was aerobically aged for at least 7 days and was then used up until the extract was 14 days old. 100 ml of this extract was added to the odor container.

Humidity is an important near-field cue for mosquitoes (Cardé and Gibson 2010; Laursen et al. 2023) and is a component of human sweat, stimuli from flowers, and oviposition sites such as those mimicked by the tested plant infusions. To determine if the relative humidity of the plume changed when sources of humidity were present in the odor container, relative humidity measurements (SHT4x sensor, Sensirion, Switzerland) were conducted, with results indicating that this aqueous odor source did not add a detectable amount of humidity to the plume (Fig. S3A). Behavioral investigations further demonstrated that humidity, in the form of 100 mL of water in the odor container, in the absence of CO2 had no effect on the recruitment rate or and its presence in combination with CO2 did not alter the spectral preference of *A. aegypti* (Fig. S3B,C).

### Visual stimuli

To gain more fine control of the spectral composition and intensity of visual stimuli we moved away from the paper targets used previously (van Breugel et al. 2015; Zhan et al. 2021; Alonso San Alberto et al. 2022; Sumner and Cardé 2022). Inspired by other systems used to generate visual stimuli for insects using LEDs (Belušič et al. 2016; Egri et al. 2020), we created a pair of LED synths to generate visual stimuli. The synths consist of an array of 17 different LEDs with peak wavelengths ranging from 390 to 743 nm (Fig. 1B). While it would have been preferable to extend the range of stimuli into the UV, we were constrained by the UV transmission of the acrylic floor of the wind tunnel (Fig. S1C). These LEDs were mounted in a hemispheric dome aimed at a central diffuse reflector made of cotton broadcloth (Fig. 1A,C). The combination of the diffuse reflection and the aperture in the top of the dome allowed us to project the light from each LED to the same circular portion of the wind tunnel floor.

The intensity of the LEDs from each of the 17 color channels in both LED synths was independently controlled using pulse width modification (PWM) using custom Arduino sketches uploaded to an Arduino Uno (Rev3, Adafruit, New York) coupled with 4 breakout boards (PCA9685, Adafruit) via a Python script (Blake and Riffell 2025). A combination of photographic and spectrographic measurements was used to tune each LED channel to produce isoquantal illumination of ∼3.5×10^11^ photons/cm2/s as measured at the surface of the wind tunnel floor. With the exception of the color intensity ramp bioassays, this intensity (1.0) was used for all monochromatic stimuli.

In order to create dark visual stimuli, it was necessary to project the light of these LED synths onto a pair of black tulle targets formed by 7 concentric circles of tulle (Fig 1C,D). These circles approximated the inverse of the Gaussian intensity distribution of the light from the LED synths. This arrangement prevented any intensity artifacts along the edge of the visual stimuli which would be present if all the tulle layers were the same diameter. These tulle targets also served to increase the saturation of colors produced by the LED synths, as the light from the synths only passed through the tulle layers once, while reflections from the fabric floor needed to pass through these layers twice (Fig. S1F).

The LED synths also allowed for the creation of composite spectra using multiple LED channels. We created achromatic gray stimuli by approximating the spectral composition of DLP projectors providing ambient illumination (Fig. S1A). This composite spectra, in combination with the tulle target, allowed us to create a neutral gray achromatic stimuli with an intensity (1.0) that closely matched the white fabric background (Fig. 1D inset). We included several controls with achromatic gray stimuli of different intensities to serve as positive and negative controls, as well as to demonstrate the effect of achromatic intensity (Fig. 2). Mid-gray stimuli with an intensity midway (0.5) between the background intensity (1.0) and the unilluminated tulle target (0.0) were used as a common control stimulus across all experiments (Figs. 2-5, S4, S5, S7).

**Figure 2.**
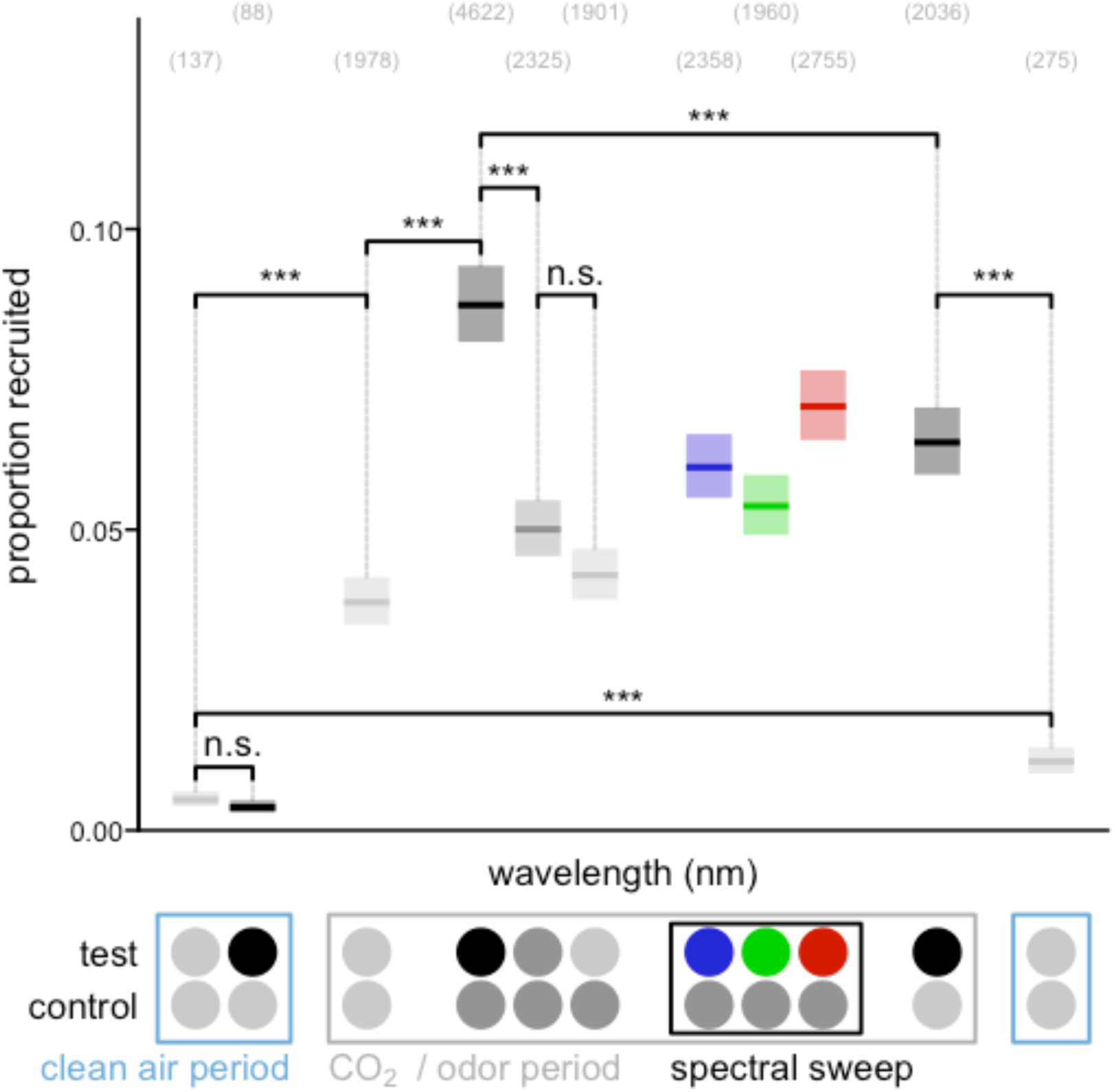
Mosquito investigations of visual stimuli are maximized by the combination of CO2 and dark visual stimuli. Proportion of trajectories investigating either the test or control visual stimulus during the spectral sweep bioassay runs. In the absence of CO2, very few mosquitoes were recruited to visual targets even if they had a low intensity (recruitment declined during the CO2 period but remained elevated relative to recruitment prior to CO2 exposure). In contrast, in the presence of CO2, mosquito recruitment was elevated to all stimuli, even those with intensity approximating the background, with darker visual targets being more preferred. Significance brackets show *post*-*hoc* contrasts with Šidák adjustment between different stimuli pairs. Test stimuli: neutral gray (light gray circles) at an intensity matching the fabric background, unilluminated black tulle targets (black circles), mid gray, and LEDs (only 435, 527 and 621 nm are shown, see Fig. S4 for full spectral sweep) at isoquantal intensities. Control stimuli: neutral or mid gray. Stimuli outside the marked CO2 / odor period were presented with clean air alone. Boxplots are the mean (line) with 95% confidence interval (shaded area). Individual points predictions are omitted here for clarity (see Fig. S4). Bracketed numbers above each bar indicate the number trajectories recruited over 100 replicate bioassay runs. Asterisks denote statistical differences: \**P* < 0.05, \*\**P* < 0.01, \*\*\**P* < 0.001

We performed two main experimental series, spectral sweeps where each of the 17 LED color channels was tested at the same intensity (Fig. 3), and color intensity ramps where we selected a limited set of LED color channels and displayed them at a range of intensities (0.00-3.00; Fig. 5). These experimental series were performed with CO2 alone and with the combination of CO2 and floral, host and oviposition site odors. In both experimental series, the order of stimuli was alternated so that stimuli appearing near the beginning of one series would appear near the end of the next series, thereby controlling for any increases or decreases in mosquito responsiveness to the visual stimuli over an experimental trial. Similar to previous wind tunnel experiments (Alonso San Alberto et al. 2022) these experimental series included a clean air only period before and after the main portion of the series (Figs. 2, S4, S5). Runs were excluded based on three independent criteria: (1) strong responses occurred to visual stimuli during these clean air-only periods (suggesting odor contamination), (2) visual responses were generally low throughout the experiment even during odor exposure (IE problems with CO2 release or experimental animals), or (3) if visual responses were much lower in one half of the series than the other (suggesting run was falling outside of the sunset activity peak).

**Figure 3.**
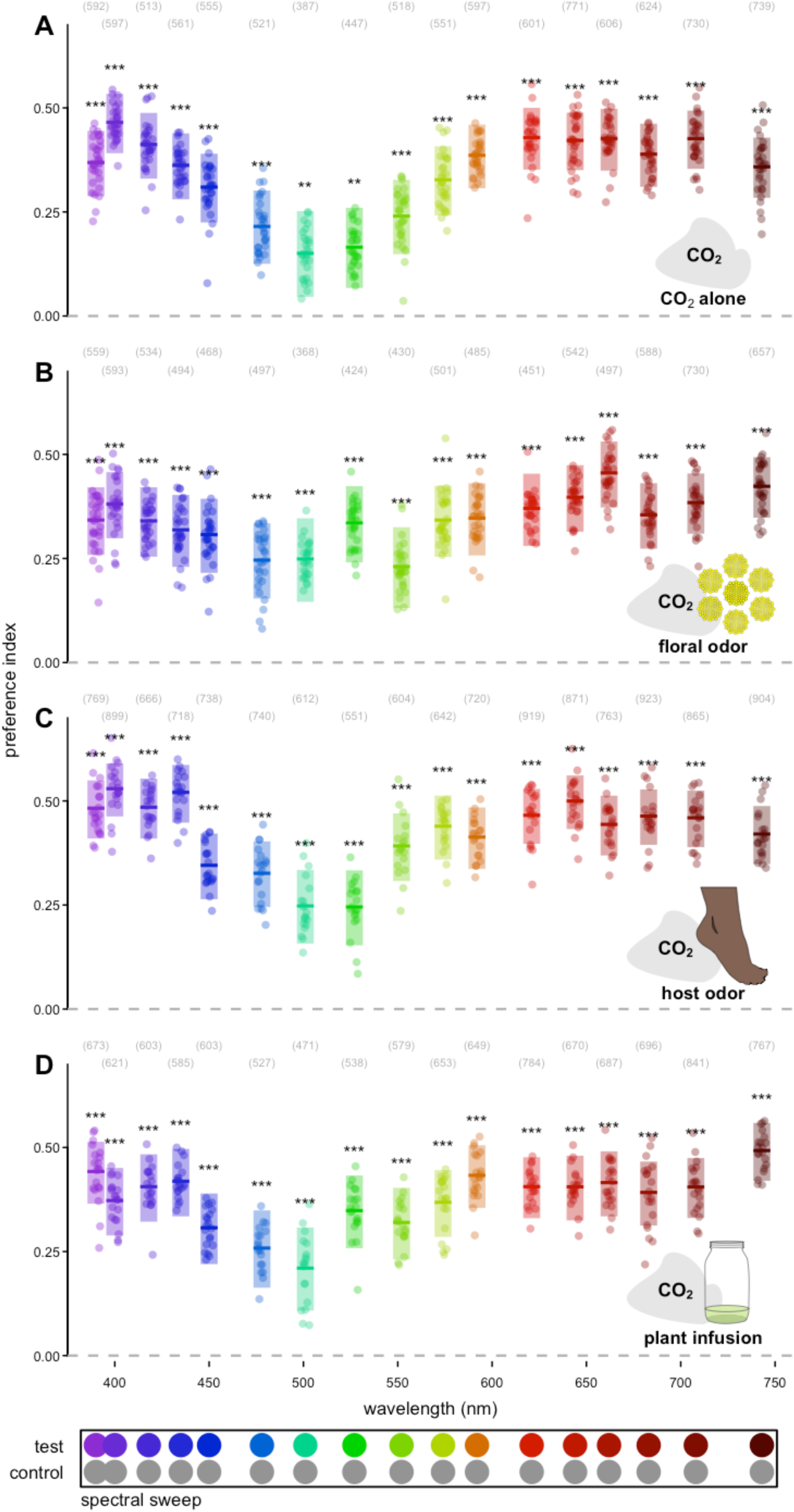
Effect of odor on mosquito spectral preference. Spectral preference in the presence of CO2 paired with (**A**) clean air, (**B**) tansy floral odor, (**C**) human foot odor, and (**D**) the odor of an alfalfa infusion. Test stimuli: LEDs at isoquantal intensities ranging from 390-743 nm. Control stimuli: mid gray. Boxplots are the mean (line) ± 95% confidence interval (shaded area), with points representing model predictions for each replicate bioassay run. Bracketed numbers above each bar indicate the number of recruited trajectories over 30, 30, 20, and 20 replicate bioassay runs. Asterisks above the boxes indicate a statistically significant difference from a preference index of 0.00. \**P* < 0.05, \*\**P* < 0.01, \*\*\**P* < 0.001

**Figure 4.**
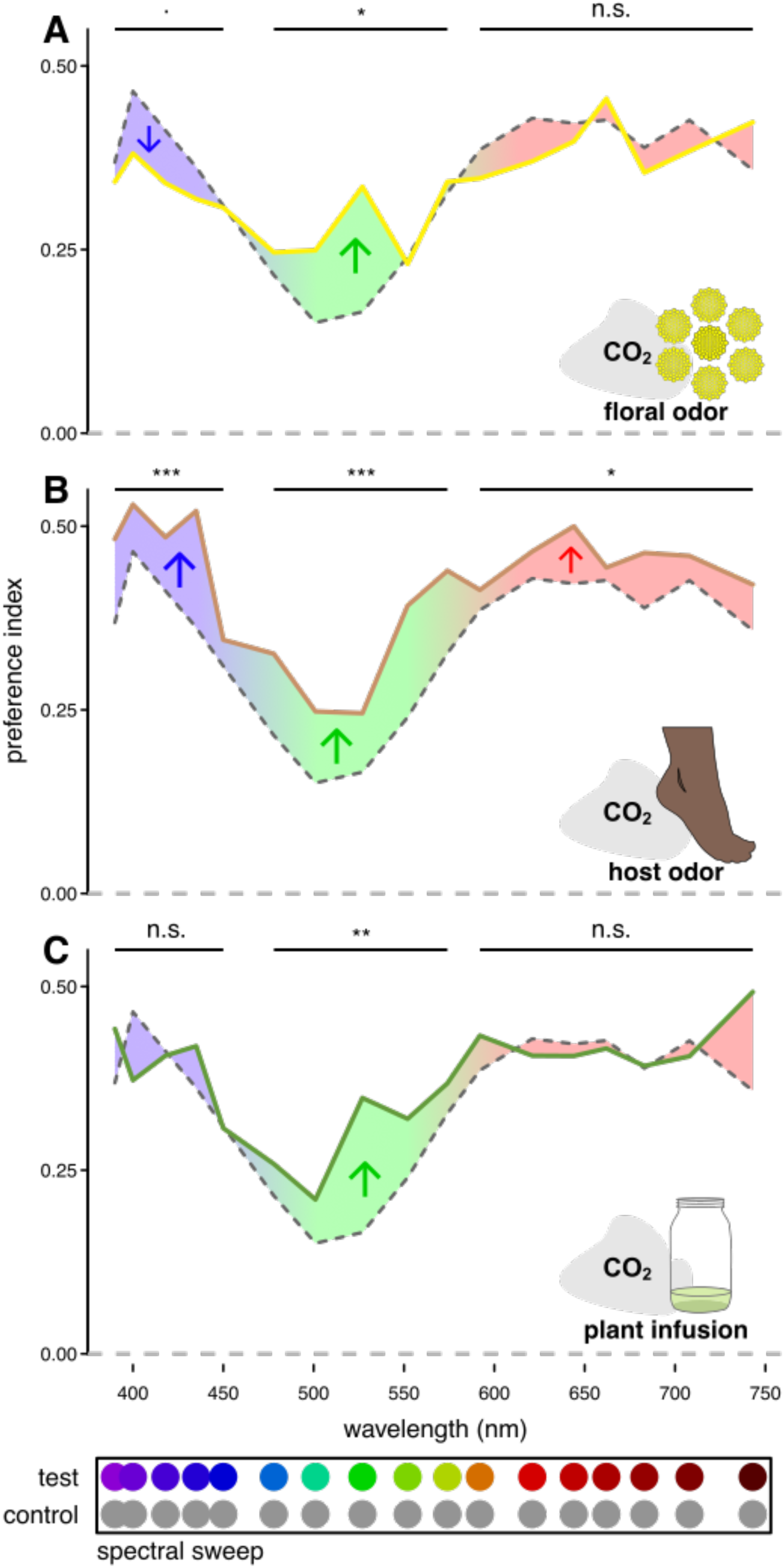
Odor shifts in spectral preference from CO2 alone. Shifts in spectral preference relative to CO2 alone (gray dashed line) with (**A**) tansy floral odor, (**B**) human foot odor, and (**C**) the odor of an alfalfa infusion. Test stimuli: LEDs at isoquantal intensities ranging from 390-743 nm. Control stimuli: mid gray. Significance stars indicate a significant difference in preference index over the specified range as determined by *a priori* contrasts. Colored arrows indicate the direction of this shift. n.s. > 0.05, ·*P* < 0.10, \**P* < 0.05, \*\**P* < 0.01, \*\*\**P* < 0.001

**Figure 5.**
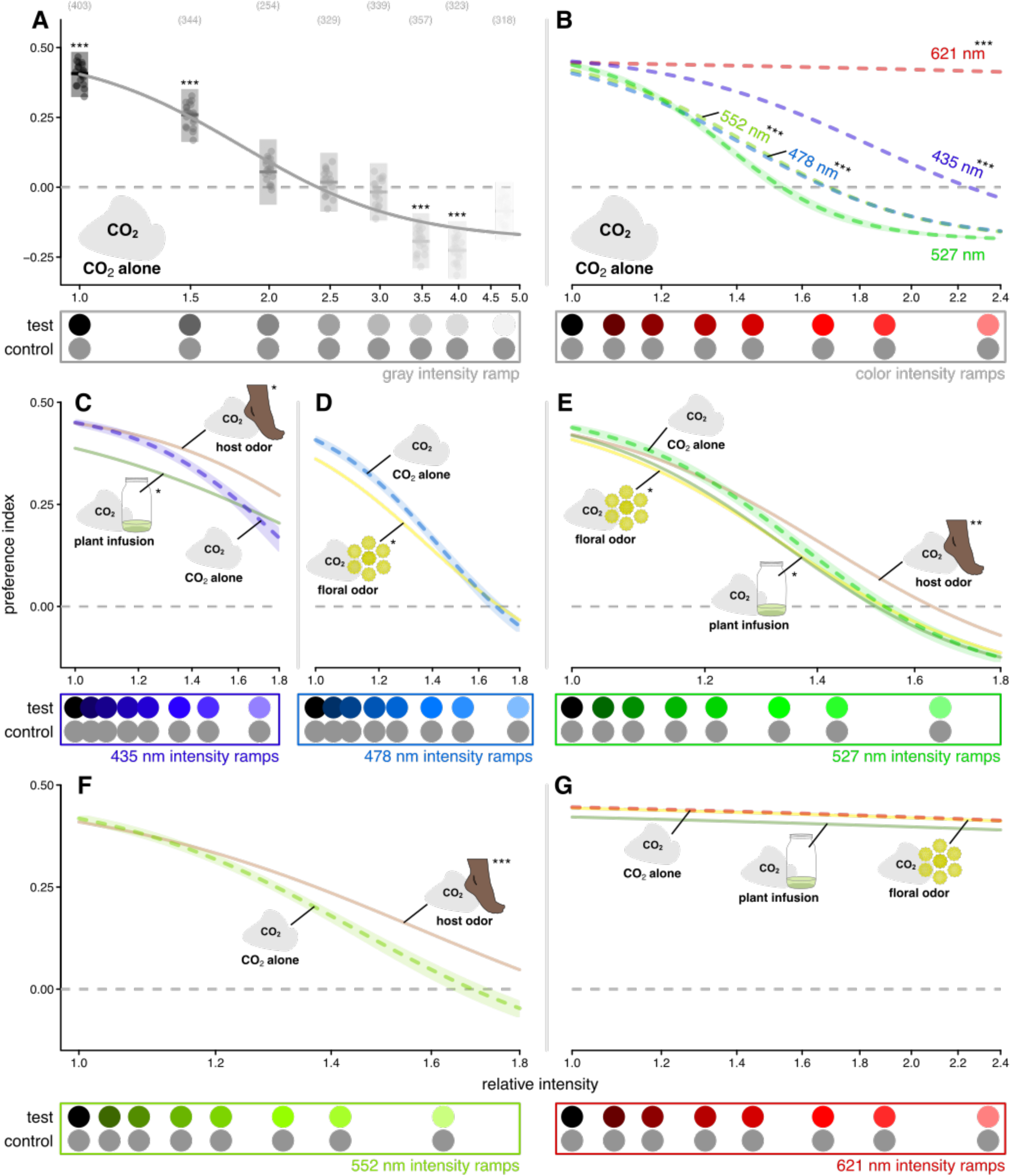
Effect of odor and wavelength on mosquito intensity preferences. We investigated the effect of intensity at a selection of wavelengths covering the visible range and focusing on spectral ranges where we observed odor-driven shifts in spectral preferences. The intensities on the x-axes are measured relative to the unilluminated black tulle targets, which was common among all of the intensity ramps, and non-zero due to the ambient illumination. Only the fitted lines are shown in **B**-**G,** graphs similar to **A** are presented in Fig S7 for all fitted lines. **A** The preference index of mosquitos responding to gray stimuli in the presence of CO2 alone. **B** The relationship between preference and relative intensity at different wavelengths as compared with the 527 nm (for clarity only the 621 intensity ramp is depicted). The effect of odor on the relationship between preference and relative intensity (not all odors were tested at all wavelengths) at (**C**) 435 nm, (**D**) 478 nm, (**E**) 527 nm, (**F**) 552 nm, and (**G**) 621 m, with all other odors compared against CO2 alone. Test stimuli: 435 nm, 470 nm, 527 nm, 552 nm, and 621 nm LED intensity ramps ranging in intensity from 0.0 to 3.0 times the isoquantal intensity used in the spectral sweep experiments, and a gray ramp ranging in intensity from 0.0 to 1.5 times the intensity of the fabric background. Control stimuli: mid gray. Boxplots in **A** are the mean (line) with 95% confidence interval (shaded area), with points representing model predictions for each replicate bioassay run. Bracketed numbers above each bar in indicate the number of recruited trajectories over 20 replicate bioassay runs. Significance stars above the boxes indicate a difference from a preference index of 0.00. Lines show a sigmoid fitted to the preference data in Fig S7. The semitransparent polygons indicate a 95% confidence interval around the line and for clarity are only displayed for the line serving as the reference for the statistical comparison. This polygon was in omitted in **G,** as due to the weak relationship with intensity the confidence interval encompassed the entire plot. Asterisks associated with a line label indicate that the relative intensity at the line’s inflection point statistically differed from that of the reference line. \**P* < 0.05, \*\**P* < 0.01, \*\*\**P* < 0.001

### Trajectories analysis

A 3D real-time tracking system (van Breugel et al. 2015; Stowers et al. 2017) was used to track the mosquitoes’ trajectories. As described above, for each experimental trial we released a group of 50 mosquitoes as this number was found to be the best compromise between minimizing interaction between individuals while still providing ample opportunities to capture mosquito responses to visual stimuli (Alonso San Alberto et al. 2022). This tracking system is unable to maintain mosquito identities for extended periods of time, however this non-independence was accounting for using mixed statistical models (see below). Previous studies have showed similar behavior between groups and singular mosquitos when responding to stimuli within this wind tunnel (Alonso San Alberto et al. 2022). Analyses were restricted to trajectories that were at least 90 frames (1.5 s) long. Occupancy maps (Fig 1F) were calculated by taking the number of mosquito occurrences within each 0.3 cm^2^ square of the wind tunnel and divided the number by the total number of occurrences in all squares yielding a percentage of residency. To evaluate the behavioral responses to the visual stimuli, three different metrics were analyzed: (1) the number of mosquito trajectories, (2) the proportions of trajectories responding to stimuli, and (3) preference indices. To assess the propensity of mosquitos to fly during a particular set of stimuli, we compared the number of recorded trajectories among stimuli, henceforth referred to as activation. We characterized mosquito preference and recruitment by trajectory passage through a pair of fictive cylinders centered on each visual stimuli (diameter: 14 cm, height: 4 cm). A sensitivity analysis demonstrated that this volume best captured the mosquitoes responding to visual stimuli (Alonso San Alberto et al. 2022). To examine the relative number of mosquitos approaching the visual stimuli, henceforth referred to as recruitment, we compared the proportion of trajectories entering either the test or control volumes. Both of these analyses were done at the stimulus level, however we evaluated preference for each trajectory (Fig. 1E) with a preference index. This index was used to evaluate and compare the preference for the test stimulus over the control stimulus irrespective of the strength of the mosquito recruitment. The index was defined as the amount of time a mosquito trajectory spent in the test stimulus volume divided by the total time it spent in both the test and control stimuli volumes. During analysis (see section below) overall preference indices were then estimated for each stimulus, weighting the contributions of trajectories by the total time spent in the stimuli volumes.

### Statistical analysis

We performed all analyses and prepared graphs using R statistical software (v4.4.1; R Core Team 2024). We analyzed the effects of visual stimulus wavelength and intensity as well as odor on mosquito preference (the proportion of time each trajectory spent in the test stimulus volumes relative to the control) using generalized mixed models (glmmTMB package, Brooks et al. 2017) with beta-binomial errors and a logit link function (Blake and Riffell 2025). We accounted for the non-independence of trajectories in the same run by including the run as a random factor, estimating individual run intercepts for each stimulus pair presented. Similar models were also examined for mosquito activation (number of mosquito trajectories during a stimulus period) and recruitment (the proportion of trajectories entering either the test or control volumes), however here the analysis was on a per run rather than a per trajectory basis. Consequently, in this case non-independence was accounted for by including a single random intercept for each run. The overall effect of stimulus factors (IE wavelength) was estimated by comparing models with and without the relevant factor using likelihood-ratio tests. Comparisons between treatment levels or groups of treatment levels were performed using the ‘eemeans’ package either as *a priori* contrasts or post-hoc contrasts with Šidák adjustment.

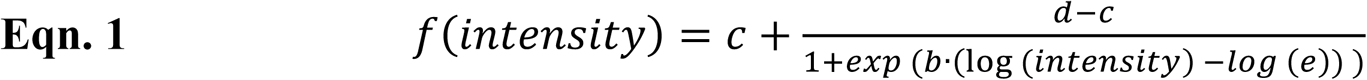

We then used the ‘medrc’ package (Gerhard and Ritz 2018) to estimate the sigmoid relationships between relative intensities of the visual stimulus and mosquito preferences (Blake and Riffell 2025). To control for the variation among wavelengths in illumination provided by the projectors (Fig. S1A), the intensities used in these models are measured relative to the black tulle target, which was common among all of the intensity ramps. We estimated individual run preference predictions for each stimulus pair presented using the mixed models detailed above and used these values as the response variable in these intensity models. These models used a four parameter log logistic model (Eqn 1), with parameters *c* and *d* representing the lower and upper asymptotes respectively, parameter *e* representing the relative intensity at the inflection point, and *b* representing the relative steepness of the curve at the inflection point. The upper and lower asymptotes were estimated from the gray intensity ramp as we saw the greatest range of relative intensity for this set of stimuli. We assumed similar upper and lower bounds to intensity preference among different wavelengths and values of *c* and *d* were held fixed for the remaining models. We estimated individual *b* and *e* values for all combinations of wavelength and odor shown in Figs. 5 and S7. We accounted for case non-independence in these models by incorporating run as a random factor, estimating *b* and *e* values for each individual run. We then used the ‘EDcomp’ function from the ‘drc’ package (Ritz et al. 2015) to test for differences in the fitted relationships by comparing the relative intensity (equivalent to relative potency) at the line’s inflection point (midway between the lower and upper bounds).

## Results

We analyzed several aspects of mosquito behavior within our behavioral bioassays. In order to characterize the effect of stimuli on activation or the propensity of mosquitos to take flight, we compared the number of trajectories recorded within a given stimulus period. Then to examine the rate at which activated mosquitos were recruited to visual stimuli, we compared the proportion of trajectories approaching either of the visual stimuli. Lastly to evaluate mosquito preferences we evaluated preferences indices, which are defined as the proportion of time each trajectory spent in the test stimulus volume relative to the control volume. These metrics in combination gave us a sense of both if mosquitos would respond to stimuli and what stimulus they would choose if they responded.

### Effect of CO2 on mosquito activation and recruitment

As CO2 is known to be a critical cue sensitizing response to visual cues (van Breugel et al. 2015; Carnaghi et al. 2021), we first wanted to confirm its effects in our bioassays. Across all experiments, we saw relatively low levels of activation in the absence of CO2 (Figs. S2,3,6) as well as very low rates of recruitment to visual cues (Figs. 2, S2-6). Less than 1% of mosquito trajectories investigating either the test or control stimulus when CO2 was absent, with most runs seeing no trajectories recruited. Recruitment was low even when highly attractive low-intensity (IE black) visual cues were used. Recruitment rates without CO2 were so low relative to CO2 rates that it made estimating preference impractical. In the presence of CO2 recruitment rates increased between 7 to 17-fold depending on the stimulus (Fig. 2). The increase in recruitment was lowest when both stimuli were gray (1.0) with an intensity closely matching the white fabric background. This increase was much greater for stimuli with a greater contrast with background. We did observe statistically significant declines in both activation (*post-hoc* contrast with Šidák adjustment, z =-13.61, *P* < 0.0001) and recruitment (*post-hoc* contrast with Šidák adjustment, z =-10.14, *P* = 0.015) over the course of the spectral sweep experiment as evidenced by the difference between the initial and final response using a unilluminated black tulle target. However, the effect of these declines was accounted for in experiments by alternating the order of stimulus presentation. We also observed that while recruitment did decline sharply after the CO2 period ended, it did not fully return to the rate seen before the CO2 release, with the post-CO2 recruitment being approximately double that occurred prerelease.

As our past work did not examine how differences in CO2 concentrations in an odor plume could affect mosquito recruitment (Alonso San Alberto et al. 2022), and the amount of CO2 varies considerably among different resources (Geier et al. 1999; Peach et al. 2019a), we evaluated the mosquito recruitment when they were provided with a plume of 0%, 1%, 5%, and 10% CO2 (Fig. S2). We found that CO2 concentration resulted in a statistically significant increase in both mosquito activation (likelihood-ratio test, χ^2^ = 125.17 df = 15 *P* < 0.0001) and recruitment (likelihood-ratio test, χ^2^ = 305.67 df = 15 *P* < 0.0001). Activation increased across all CO2 concentrations but had a relatively high baseline as compared with recruitment. Recruitment also differed across all CO2 concentrations, with the increase beginning to plateau at 5%. The effect of CO2 was also smaller for gray stimuli with minimal contrast with the fabric background.

Despite the strong effect of CO2 on recruitment, we did not observe any statistically differences in preference for visual stimuli among the different CO2 concentrations (likelihood-ratio test, χ^2^ = 12.55 df = 10 *P* = 0.25) and they were largely consistent with the spectral and intensity preferences observed in the spectral sweep and intensity ramp experiments.

### Effect of wavelength and intensity on mosquito activation and recruitment

We also examined the effects of wavelength and intensity on mosquito activation and the rates of mosquito recruitment (Fig. S2, S4, S5). While stimulus wavelength (likelihood-ratio test, χ^2^ = 58.69 df = 64 *P* = 0.66) and intensity (likelihood-ratio test, χ^2^ = 115.78 df = 105 *P* = 0.22) did not statistically significantly affect mosquito activation, both wavelength (likelihood-ratio test, χ^2^ = 164.36 df = 64 *P* < 0.0001) and intensity (likelihood-ratio test, χ^2^ = 224.79 df = 105 *P* < 0.0001) had a significant effect on recruitment in the spectral sweep (Fig. S4) and intensity ramp experiments (Fig. S5) respectively. Stimuli with high intensity or stimuli at or near the green range (500-550 nm) where mosquitos have higher spectral sensitivity resulted in decreased mosquito recruitment. However, these effects on recruitment were small compared to the effects on visual preference (compare Figs. 4,5).

### Effect of odor and humidity on mosquito activation and recruitment

In the spectral sweep, we found that the odor of human feet and alfalfa plant extract significantly increased both mosquito activation (likelihood-ratio test, χ^2^ = 75.06 df = 51 *P* = 0.0158) and recruitment to visual stimuli (Fig. S4, likelihood-ratio test, χ^2^ = 87.76 df = 51 *P* = 0.0010).

However in the intensity ramp experiments, we did not observe a similar increase with odor in either activation (likelihood-ratio test, χ^2^ = 87.38 df = 72 *P* = 0.10) or recruitment (Fig. S5, likelihood-ratio test, χ^2^ = 81.83 df = 72 *P* = 0.20). As it was operationally complicated to alternate between odors, different odors were tested at different times. Therefore, it is probable that these differences in recruitment observed during the spectral sweep experiments could be attributed to differences in responsiveness among the different cohorts of experimental animals.

To better evaluate the effect of odor on mosquito recruitment, we performed an additional experiment where we tested foot odor concurrently with and without CO2. In this experiment, we found the addition of foot odor to as compared with CO2 alone resulted in a much smaller though statistically significant increase in recruitment (Fig. S6, *a priori* contrast, z =-2.48 *P* = 0.0129) as compared with the increase seen in the spectral sweep and intensity ramp experiments.

Additionally there was a small but statistically significant increase in recruitment with foot odor over filter air in the absence of CO2 (*a priori* contrast, z =-2.42, *P* = 0.0153). Foot odor also decreased mosquito activation when CO2 was present (*a priori* contrast, z = 3.34, *P* = 0.0008).

We also performed a similar experiment to examine the potential confounded effects of humidity produced by some of our odor sources. Despite humidity being an import host cue (Cardé and Gibson 2010), we did not observe an increase in activation or recruitment when humidity was presented in the absence of CO2 (Fig. S3B,C). This lack of response might be due to the low amount of humidity (Fig. S3A), or that the humidity was not co-localized with the visual cues and humidity is a short range cue (Laursen et al. 2023). However in the presence of CO2 humidity actually decreased both activation (*a priori* contrast, z = 2.16, *P* = 0.0310) and recruitment (*a priori* contrast, z = 3.70, *P* = 0.0002), perhaps due to more mosquitos being attracted to the odor outlet (and source of humidity) rather than to the visual stimuli. We also did not detect any statistically significant differences in spectral preferences with humidity as compared with CO2 alone (Fig. S3D).

### Effect of odor on mosquito spectral preference

Our prior investigation of mosquito spectral preferences used paper targets with broad spectral composition and was unable to fully control for differences in intensity (Alonso San Alberto et al. 2022). Refining those earlier methods, we used narrowband isoquantal stimuli generated by a pair of LED synths. Unlike the other experiments, these spectral sweep experiments had stimuli spanning a wide spectral range allowing us to better characterize spectral preference. Preference in this and other experiments was defined as the proportion of time each trajectory spent in the test stimulus volume relative to the control volume. We found that preference measurements were more consistent across cohorts as compared with recruitment (compare Figs. 3, S7 to Figs. S4, S5)

In the presence of CO2 alone, mosquito exhibited preferences for shorter wavelengths (390 nm – 420 nm), with the preference declining gradually as the stimulus wavelength approached the green range (500-550 nm), before increasing and subsequently plateauing above 600 nm (Fig. 3). Despite a decreased preference for stimuli in the green range (preference index of ∼ 0.15), all the examined colored stimuli in the spectral sweep were more attractive than the moderately attractive medium grey control. Outside the green range we saw preference indices approaching 0.50, similar to the attraction we observed for the highly attractive unlit black tulle target (Fig. 5A).

In contrast to our previous work that only tested CO2, here we conducted spectral sweep experiments with different olfactory stimuli. We found that odors other than CO2 resulted in statistically significant differences in the spectral preferences of *Ae. aegypti* (Fig. 4, likelihood-ratio test, χ^2^ = 93.62 df = 51 *P* = 0.0003). Floral odors resulted in a non-significant negative preference shift in the lower wavelength range (390-450 nm), and a positive shift in the green range (475-575 nm). The odor of a plant infusion (oviposition lure) also increased the attraction in the green range, but this shift appears to be more towards the longer green wavelengths than with floral odor. There was also no shift in in preference in the short wavelength range with the plant infusion odor. In contrast to the other odors, foot odor generally increased the preference to all stimuli, with statistically significant shift in all wavelength ranges (Fig. 4).

### Effect of odor on mosquito intensity preferences

As has been noted previously (van Breugel et al. 2015; Alonso San Alberto et al. 2022), the preference of mosquitos for a visual stimulus is highly dependent on its intensity (Fig. 5A), with darker stimuli being highly preferred. We also found that the effect of intensity of mosquito preference was wavelength dependent, with the sharpest declines with increased intensity seen with the 527 nm stimuli (Fig. 5B). Stimuli with wavelengths further from the green showed more shallow declines, with the 625 nm stimuli showing no appreciable decline with increased intensity. We saw significant flattening of these intensity relationships with the addition of foot odor (Fig. 5C,E,F). In contrast, floral odor and alfalfa extract had comparably small effects on these intensity relationships (Fig. 5C,D,E,G) and in some cases these odors even had a marginal negative effect on preference (Fig. 5D,E).

## Discussion

Previous behavioral investigations of vertebrate host finding in mosquitos have demonstrated that this process involves the integration of olfactory, visual, thermal, and tactile cues. Color or more specifically the spectral composition of visual cues has also been shown to be an important during host finding. This study builds on this previous work by investigating the effects of odors beyond CO2 on the spectral preferences of *Ae. aegypti*. These odors suggestive of vertebrate hosts, floral resources, and oviposition sites all resulted in shifts in the spectral preferences of mosquitos, suggesting these color preferences are context dependent.

### Mosquito spectral preferences

Consistent with long-standing observations that mosquitos are attracted to dark (IE black) high contrast visual stimuli during vertebrate host-search (Kennedy 1940; Sippell and Brown 1953; Muir et al. 1992a; van Breugel et al. 2015), floral foraging (Peach et al. 2019b), and during oviposition (Snow 1971), we found that *Ae. aegypti* females showed a strong preference for similar dark stimuli (Fig. 5). We also found that similar to previous studies (Brown 1954; Muir et al. 1992a; Alonso San Alberto et al. 2022), mosquitos preferred longer wavelength colors (> 580 nm, orange, red) over shorter wavelengths (< 570 nm). However, unlike previous studies the use of our LED generated stimuli as compared with fabric or paper targets allowed us to disentangle a stimuli’s spectral content or chromaticity from its intensity and contrast with the background. This decoupling better reveals an inverse gaussian shaped curve of preference centered in the green with decreased attraction to both blue and violet as well as red and oranges colors (Fig. 3A). Our past work (Alonso San Alberto et al. 2022) observed a preference for cyan wavelengths (470-510 nm) that was not observed in this study. This difference with our previous study may be accounted for by the narrow spectral range of the LED stimuli with an intensity more closely matching the other colors. As shown in Fig. 5B, mosquitoes are maximally sensitive (at least among the tested wavelengths) to green wavelengths, as it was with these wavelengths where we observed the steepest declines with increased intensities. This result mirrors the action spectrum shown in ovipositing *Ae. aegypti* (Snow 1971), where green light had the greatest effect on oviposition preference. Our results suggest that the decreased preferences to green reflect the innate preference for dark low intensity stimuli, and do not demonstrate color discrimination.

While the responses of *Ae. aegypti* to red wavelengths (Fig 5B,G) showed no evidence of sensitivity to light beyond 600 nm, this result does not preclude such sensitivity as we tested only a relatively narrow range of intensities. As the sensitivity of opsins decreases in an exponential manner as you move away from the peak sensitivity (Stavenga et al. 1993), we would expect residual sensitivity to extend well into the 600-700 nm range. At higher intensities optomotor responses have been demonstrated in *An. gambiae* with wavelength >600 nm (Gibson 1995).

Female *Ae. aegypti* have also been shown to discriminate between green and red stimuli of matched intensities further suggesting sensitivity to wavelength beyond 600 nm (Alonso San Alberto et al. 2022).

### Olfactory gating of visual preferences

This study is the first to examine how odors beyond CO2 can gate and change the visual preferences of mosquitos. Mosquitos in different behavioral contexts are searching for different resources (vertebrate hosts, nectar sources, oviposition sites), and the visual appearance of these resources can differ greatly (Alonso San Alberto et al. 2022). It has been suggested that odor could gate responses to other resources in a way similar to the way CO2 gates the response to vertebrate host cues (Alonso San Alberto et al. 2022; Shannon et al. 2024). In this study we found that odors associated both with attractive floral resources (Peach et al. 2019a) and oviposition sites (Ritchie et al. 2014) significantly increased mosquito preference for green (475-

575 nm) stimuli (Fig. 4). These responses were also gated by CO2, but we did find that concentrations similar to those possibly emitted by flowers (Peach et al. 2019a) were sufficient to activate visual responses (Fig. S2). Many flowers including the common tansy used in our experiments have a reflectively high reflectance in this range (Arnold et al. 2010; Peach et al. 2019b), and an increased to this range of wavelengths would be expected to increase floral foraging success. The increase in sensitivity to green wavelengths for the oviposition lure is less clear but might indicate water enriched with plant matter. In contrast with the other odors, human foot odor increased visual response across the entire range of tested wavelengths (Fig. 4). This result is unsurprising as host odor synergizes with CO2 to increase mosquito response to visual stimuli (Lacey et al. 2014; Cardé 2015).

Coupling between the olfactory and visual systems in mosquitos has already been demonstrated with CO2 stimulation greatly increasing both visual tracking (Barredo et al. 2022) and physiological responses in the lobula (Vinauger et al. 2019). In Diptera this visual neuropil responds to moving objects, seems to have a role in target detection and relays chromatic information to the central brain (Nordström and O’Carroll 2006; Trischler et al. 2007; Aptekar et al. 2015; Lin et al. 2016). In contrast visual stimuli do not seem to modulate responses within the antennal lobe, where olfactory information is processed in mosquito brains (Vinauger et al. 2019). Given this established pathway between these two processing centers, it seems likely that other odors could similarly modulate mosquito visual responses. It has been hypothesized that the low acuity of mosquito vision might preclude object identification, leaving odor as the primary source of information (Vinauger et al. 2019).

### Olfaction differently moderates responses to chromatic and achromatic cues

From the observed spectral preferences (Fig. 3A), it seems likely that in the presence of CO2 alone *Ae. aegypti* visual response is dominated by achromatic contrast cues, with the level of contrast being variable across the mosquito visual range due to differences in spectral sensitivity. The observed spectral preferences are inversely proportional to the electroretinogram-determined spectral sensitivity of the *Ae. aegypti* compound eye, with a primary sensitivity peak in the green at ∼525 nm and a secondary peak in the UV ∼ 350 nm (Muir et al. 1992b). Like in other Dipterans, the R1-6 outer photoreceptors are the most numerous type in mosquitos, and these photoreceptors dominate electroretinogram responses (Minke et al. 1975). While these photoreceptors express a green-sensitive opsin (Hu et al. 2012), it seems likely that they also express a UV-sensitizing pigment similar to the outer photoreceptors in Brachyceran flies, giving them UV sensitivity (Stavenga et al. 2017). Given this broadened sensitivity, the visual response of mosquitos to targets could be explained through the responses of R1-6 photoreceptors alone. However, it seems that a subset of the R7 central photoreceptors may be sufficient if the function of the outer photoreceptors is disrupted (Zhan et al. 2021).

While the preferences of mosquitos with CO2, alone seem dominated by the responses of R1-6 photoreceptors, the shifts in spectral preference with other odors suggest that this general preference for dark contrasting stimuli is modified by input from the central photoreceptors. The opsin expression in *Ae. aegypti* central photoreceptors show that R8 cells throughout most of the eye express a green-sensitive opsin (Hu et al. 2012), with R7 cells co-expressing either a green and blue sensitive option in the dorsal and ventral stripe regions or a green and UV sensitive option in the remainder of the eye (Hu et al. 2009; Hu et al. 2011; Hu et al. 2014). In *Cx. pipiens*, electroretinogram with chromatic adaptation suggests the presence of photoreceptors primarily sensitive to green and UV wavelengths (Peach et al. 2019b). Taken together, this suggests that mosquitos could discriminate by comparing the inputs of R7 and R8 cells in an opponent matter similar to that seen in *Drosophila* (Schnaitmann et al. 2020). This opponency could allow green wavelengths to have a positive input on spectral preference that modifies but not negates the preference for dark objects. Our results suggest that this input is gated by odor and not a general aspect of mosquito visual response.

Despite the observed shift in preference during the spectral sweep experiments (Fig. 4), we did not observe a flattening of the intensity preference relationship in the intensity ramp experiments (Fig. 5E). It may be that the strong general preference for dark objects is masking the effect in these experiments, and this spectral discrimination could be more apparent when mosquitos are presented with a choice between two differently colored quasi-monochromatic stimuli rather than a choice between one such stimulus and an achromatic control. It is also possible that size of this spectral discrimination could be modulated by physiological state. For example, younger mosquitos are more likely to seek out floral resources, and older mosquitos are more predisposed to blood-feeding (Foster and Takken 2004; Peach et al. 2019a). We elected to eliminate physiological state as a source of variation by testing only 7 day old non-blood fed females. We would have expected younger and gravid mosquitos to respond more strongly to the floral (Peach et al. 2019a) and plant extract oviposition lures (Ritchie et al. 2014), respectively.

### Implications for control interventions

Traps integrating visual and olfactory cues have been demonstrated to be improve the capture of hematophagous insects such as tsetse flies and kissing bugs (Green 1986; Reisenman et al. 2000; Torr and Vale 2015). Visual cues are also an important aspect of mosquito trap design (Bidlingmayer 1994; Kline 2006), however the interplay of vision and olfaction is not generally considered. In addition, mosquito trap design has been largely static with little change over the last 50 years and unattractive colors (white, green, blue) are often employed (Bidlingmayer 1994; Kline 2006). The shifts in spectral preferences we observed with odor suggest that odor and visual cues in traps could be better tailored to target different species or different groups of mosquitos (IE gravid mosquitos, newly emerged mosquitos). There remains considerable work to be accomplished to refine mosquito trap design to make better use of both color and odor.

### Conclusions and future research

Building on work on olfactory gating of visual cues in mosquitos (Vinauger et al. 2019; Alonso San Alberto et al. 2022), the study demonstrated that odors other than CO2 can shift the visual responses of *Ae. aegypti*. The shifts in preference to these odors suggest that different odor cues could gate visual response in a context dependent manner, priming mosquitos to respond to differ resources in their environment. Our results further suggest that only the outer photoreceptors relating to achromatic responses are involved in responses to visual cues in the presence of CO2, however the addition of other odors can recruit input from the central photoreceptors. We only examined a limited set of odor extracts; future research should examine a wider set of odors and dissect these to discover which odorants are driving the observed shifts. As the gating of visual responses by CO2 differed among species (Alonso San Alberto et al. 2022), and mosquitos vary greatly in their ecology and host preferences, it seems likely that olfactory gating differs among mosquito species and could offer many potential avenues for study. The study of visual responses in mosquitos and their modulation by olfaction would also be aided by a deeper and more mechanistic understanding of their photoreceptors and the potential intensity, polarization (Bernáth and Meyer-Rochow 2016), and chromatic channels available as inputs for mosquito behavior.

## Acknowledgements

We would like to thank N. Salimi and K. Hanssen for assistance with behavioral bioassays, and Binh Nguyen for mosquito rearing and care work. We are also grateful for the discussions and advice provided by G. Van Susteren, C. Ruiz, G. Tauxe, F. Chen, and O. Akbari.

## Competing interests

The funders had no role in study design, data collection and analysis, decision to publish, or manuscript preparation.

## Author contributions

Conceptualization: AJB, JAR; Methodology: AJB; Software: AJB; Formal analysis: AJB; Investigation: AJB; Data curation: AJB; Writing - original draft: AJB; Writing - review & editing: AJB, JAR; Visualization: AJB; Supervision: JAR; Project administration: JAR; Funding acquisition: JAR

## Funding

Support for this project was funded by the Air Force Office of Scientific Research under grants (FA9550-20-1-0422 and FA9550-22-1-0124); National Institutes of Health under grants R01AI175152 and R01AI148300 (JAR); and an Endowed Professorship for Excellence in Biology (JAR).

## Data availability

Data and code are available from Mendeley Data: https://doi.org/10.17632/fdr7znz5dh.1 (Blake and Riffell 2025).

---

**Figure S1.**
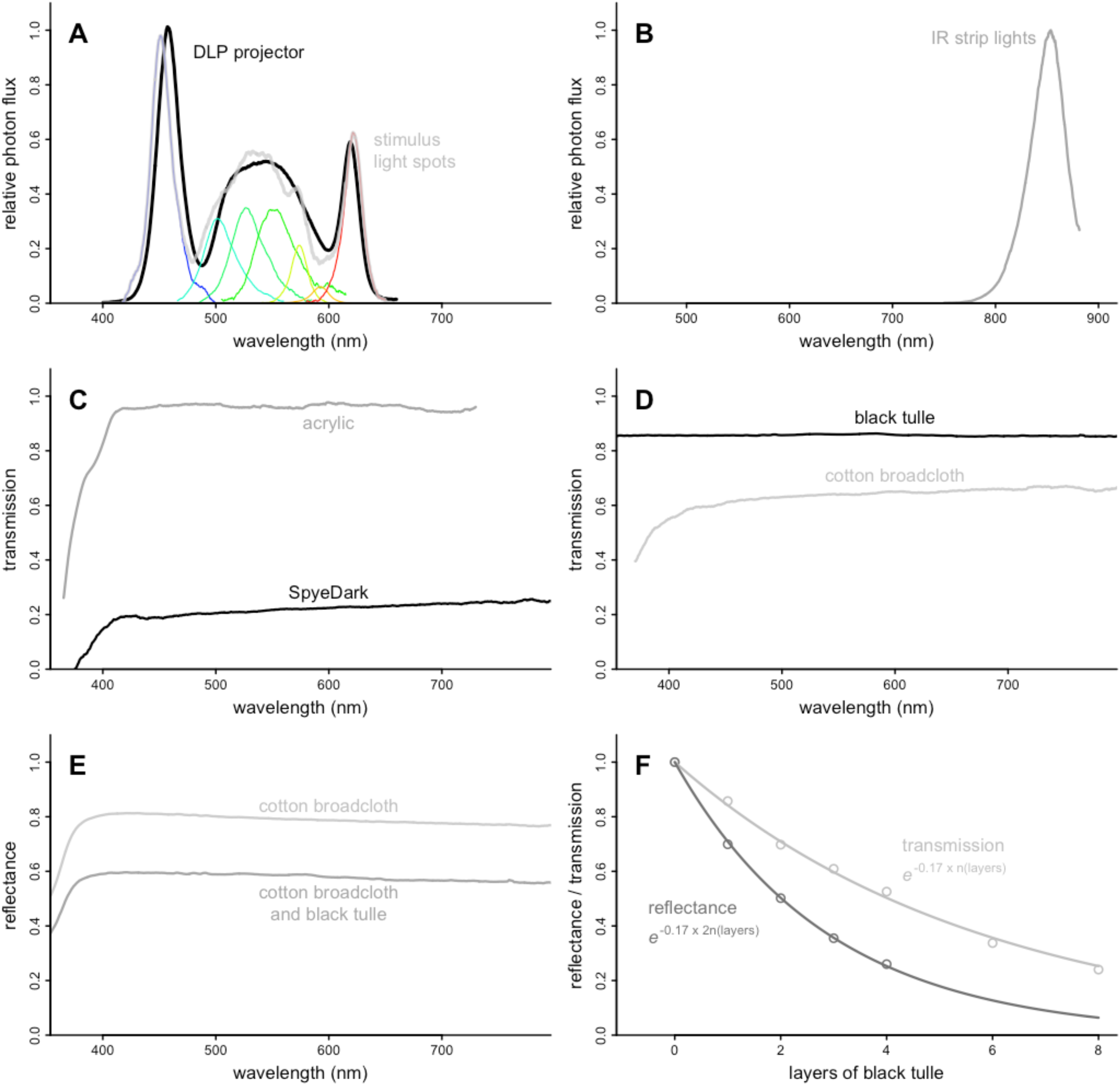
**Spectroscopy of surfaces and light sources used in the wind tunnel**. **A** Relative photon flux of the DLP projectors providing ambient illumination in the wind tunnel and of the grayscale visual stimuli. The photon fluxes of the 7 component LED color channels are shown with the thinner colored lines below. **B** Relative photon flux of the IR LED strips used to provide illumination for the cameras. **C** Transmission of acrylic walls and floor of the wind tunnel, and the SpyDark panels lining the walls. **D,E** Transmission and reflectance of the materials making up the fabric liner for the wind tunnel floor. The reflection of the cotton broadcloth and black tulle was estimated from the reflectance of cotton broadcloth corrected for transmission through two layers of black tulle (see S1f). **F** The relationship between transmission/reflectance and the number of tulle layers. Transmission measurements (light gray circles) estimated spectrographically, and reflectance measurements (dark gray circles) estimated photographically. Reflectance through a layer of tulle can be viewed as transmission through the layer by both the incident and reflected light.

**Figure S2.**
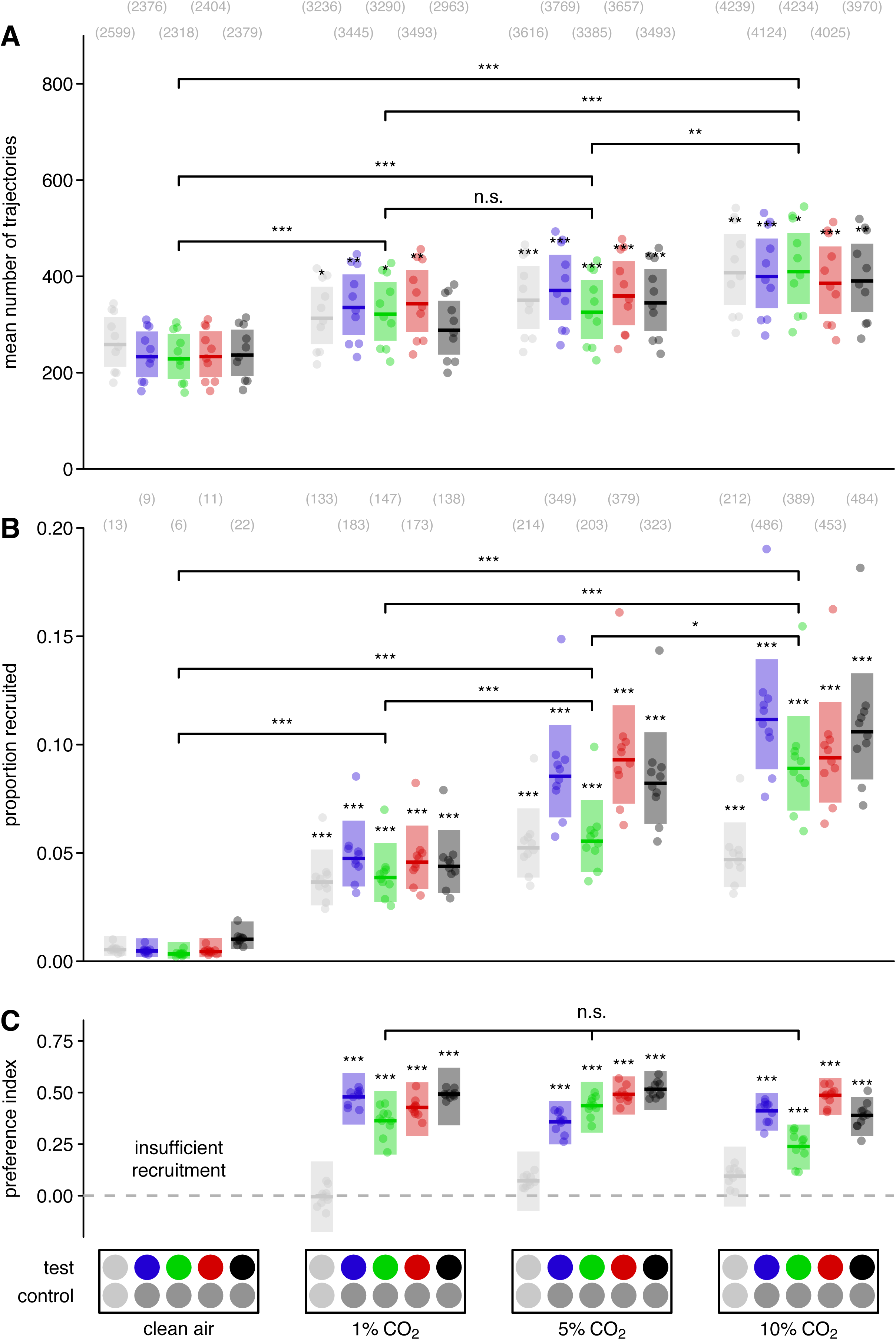
Effect of CO2 concentration on mosquito behavior. **A** The mean number of mosquito trajectories recorded, which measured mosquito activation, during each stimulus period where the plume consisted of clean air alone, 1% CO2, 5% CO2, or 10% CO2. Asterisks above the boxes here indicate a statistical difference in the number of trajectories as compared with the stimulus period with paired neutral gray stimuli and clean air alone (leftmost box). We found statistically significant differences in the number of trajectories among different concentrations of CO2 in the plume (significance brackets, *post*-*hoc* contrasts with Šidák adjustment). Bracketed numbers above each boxplot indicate the total number of trajectories over 10 bioassay runs. **B** The proportion of trajectories recruited to either the test or control visual stimuli under the same odor conditions listed above. Asterisks above the boxes here indicate a statistical difference from the recruitment to paired neutral gray stimuli with clean air alone (leftmost box). We found statistically significant differences in the recruitment to visual cues among different CO2 concentrations (significance brackets, *post*-*hoc* contrasts with Šidák adjustment). Bracketed numbers above each boxplot indicate the total number of recruited trajectories over 10 bioassay runs. **C** The preference indices of mosquitos in the wind tunnel responding to visual stimuli of various colors. Asterisks above the boxes indicate a statistical difference from a preference index of 0.00. We found no statistically significant differences in the visual preference of responding mosquitos among the 1%, 5%, and 10% CO2 concentrations (significance bracket, likelihood-ratio test, χ^2^ = 12.55 df = 10 *P* = 0.25). Test stimuli from left to right: neutral gray (light gray circles) at an intensity matching the fabric background, blue (450 nm), green (527 nm), red (621 nm) LEDs at matching isoquantal intensities, and unilluminated black tulle targets (black circles). Control stimuli: neutral or mid gray. Boxplots are the mean (line) with 95% confidence interval (shaded area), with points representing model predictions for each of replicate bioassay run. n.s. > 0.05, \**P* < 0.05, \*\**P* < 0.01, \*\*\**P* < 0.001

**Figure S3.**
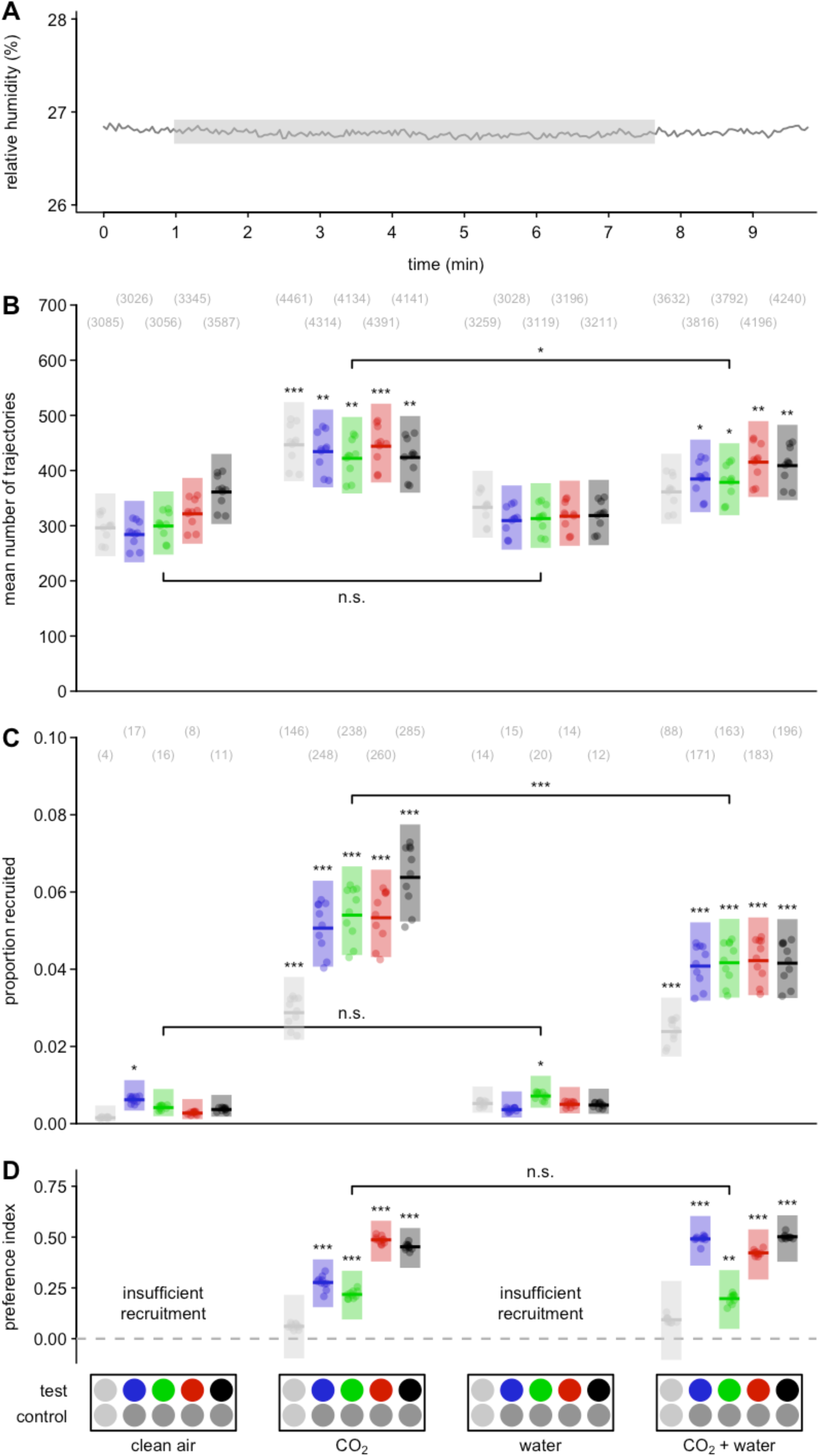
Effect of humidity on mosquito behavior. **A** Relative humidity in the wind tunnel as measured by a sensor (SHT4x, Sensirion) located 10 cm downwind of the odor/CO2 outlet. The gray box indicates the period where 10% of the plume air passed through the odor jar containing 100 ml of distilled water (humidified air), however no increase in humidity is apparent. **B** The mean number of mosquito trajectories recorded, which measured mosquito activation, during each stimulus period where the plume consisted of clean air alone, 10% CO2, humidified air, and the combination of CO2 and humidified air. Asterisks above the boxes here indicate a statistical difference in the number of trajectories as compared with the stimulus period with paired neutral gray stimuli and clean air alone (leftmost box). When CO2 was present, we found humidified air caused a significant decrease in the number of trajectories (significance bracket, *a priori* contrast, z = 2.16, *P* = 0.0310). Bracketed numbers above each boxplot indicate the total number of trajectories over 10 bioassay runs. **C** The proportion of trajectories recruited to either the test or control visual stimuli under the same odor conditions listed above. Asterisks above the boxes here indicate a statistical difference from the recruitment to paired neutral gray stimuli with clean air alone (leftmost box). When CO2 was present, we found humidified air caused a significant decrease in recruitment (significance bracket, *a priori* contrast, z = 3.70, *P* = 0.0002). Bracketed numbers above each boxplot indicate the total number of recruited trajectories over 10 bioassay runs **D** The preference index of mosquitos in the wind tunnel responding to visual stimuli of various wavelengths under the same odor conditions listed above. Asterisks above the boxes here indicate a statistical difference from a preference index of 0.00. We found no evidence that this humidified air had any effect of the visual preference of responding mosquitos (significance bracket, likelihood-ratio test, χ^2^ = 6.96 df = 5 *P* = 0.22). Test stimuli from left to right: neutral gray (light gray circles) at an intensity matching the fabric background, blue (450 nm), green (527 nm), red (621 nm) LEDs at matching isoquantal intensities, and unilluminated black tulle targets (black circles). Control stimuli: neutral or mid gray. Boxplots are the mean (line) with 95% confidence interval (shaded area), with points representing model predictions for each of replicate bioassay run. n.s. > 0.05, \**P* < 0.05, \*\**P* < 0.01, \*\*\**P* < 0.001

**Figure S4.**
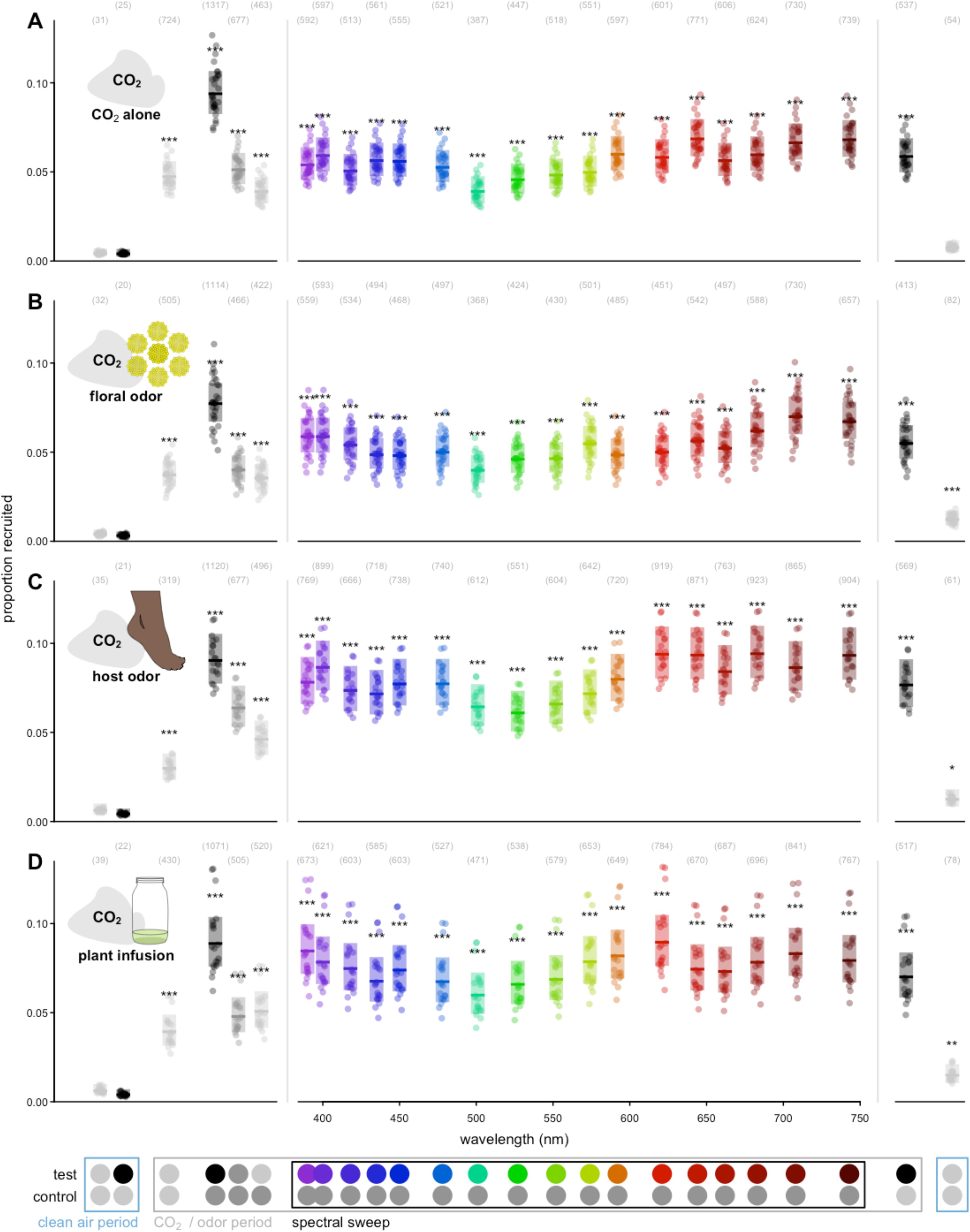
Effect of odor on mosquito recruitment to visual stimuli during the spectral sweeps. Proportion of trajectories recruited to either the test or control visual stimulus in the presence of CO2 paired with (**A**) no odor, (**B**) tansy (*T. vulgare*) floral odor, (**C**) human foot odor, and (**D**) the odor of an alfalfa infusion. Test stimuli: neutral gray (light gray circles) at an intensity matching the fabric background, unilluminated black tulle targets (black circles), and LEDs at isoquantal intensities ranging from 390-743 nm (Fig. 1e). Control stimuli: neutral or mid gray. Stimuli outside the marked CO2 / odor period were presented with clean air alone. The order of the spectral sweep stimuli was alternated between bioassay runs, with all other stimuli pairs always appearing in the order depicted above. Boxplots are the mean (line) with 95% confidence interval (shaded area), with points representing model predictions for each of replicate bioassay run. Bracketed numbers above each bar indicate the number of recruited trajectories over 30, 30, 20, and 20 replicate bioassay runs. Asterisks above the boxes indicate a statistical difference from the recruitment to paired neutral gray stimuli with clean air alone (leftmost box). \**P* < 0.05, \*\**P* < 0.01, \*\*\**P* < 0.001

**Figure S5.**
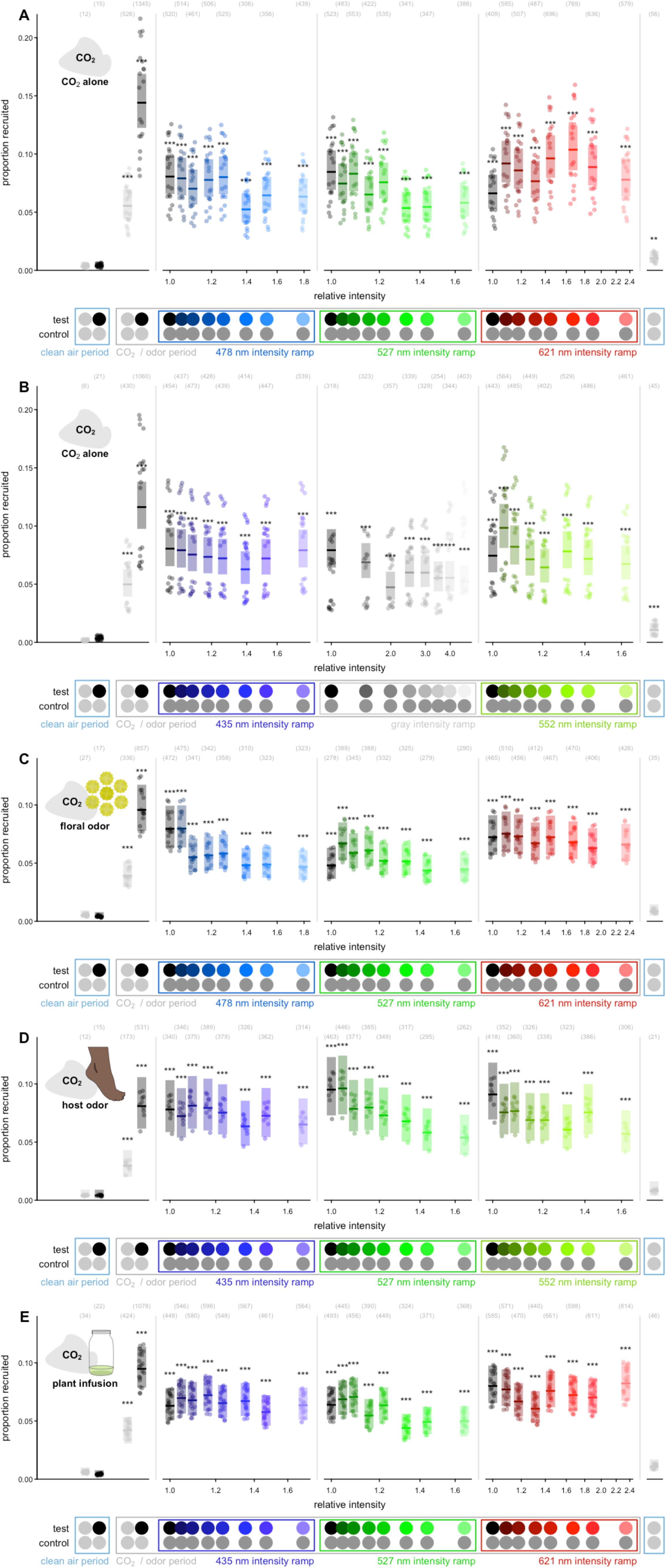
**Effect of odor and stimulus intensity on mosquito recruitment to visual stimuli**. Proportion of trajectories recruited to either the test or control visual stimulus in the presence of CO2 paired with (**A,B**) no odor, (**C**) tansy (*T. vulgare*) floral odor, (**D**) human foot odor, and (**E**) the odor of an alfalfa infusion. We investigated the effect of intensity at a selection of wavelengths covering the visible range and focusing on spectral ranges where we observed odor shifts in spectral preferences in the spectral sweep experiments. The intensities on the x-axes are measured relative to the unilluminated black tulle targets, which were common among all of the intensity ramps, and non-zero due to the ambient illumination. Test stimuli: neutral gray (light gray circles) at an intensity matching the fabric background, unilluminated black tulle targets (black circles), 435 nm, 470 nm, 527 nm, 552 nm, and 621 nm LED intensity ramps ranging in intensity from 0.0 to 3.0 times the isoquantal intensity used in the spectral sweep experiments, and a gray ramp ranging in intensity from 0.0 to 1.5 times the intensity of the fabric background. Control stimuli: neutral or mid gray. Stimuli outside the marked CO2 / odor period were presented with clean air alone. Boxplots are the mean (line) with 95% confidence interval (shaded area), with points representing model predictions for each of replicate bioassay run. Bracketed numbers above each bar indicate the number of recruited trajectories over 20, 20, 16, 10 and 20 replicate bioassay runs, respectively. Asterisks above the boxes denote a statistically significant difference from the recruitment to paired neutral gray stimuli with clean air alone (leftmost box). \**P* < 0.05, \*\**P* < 0.01, \*\*\**P* < 0.001

**Figure S6.**
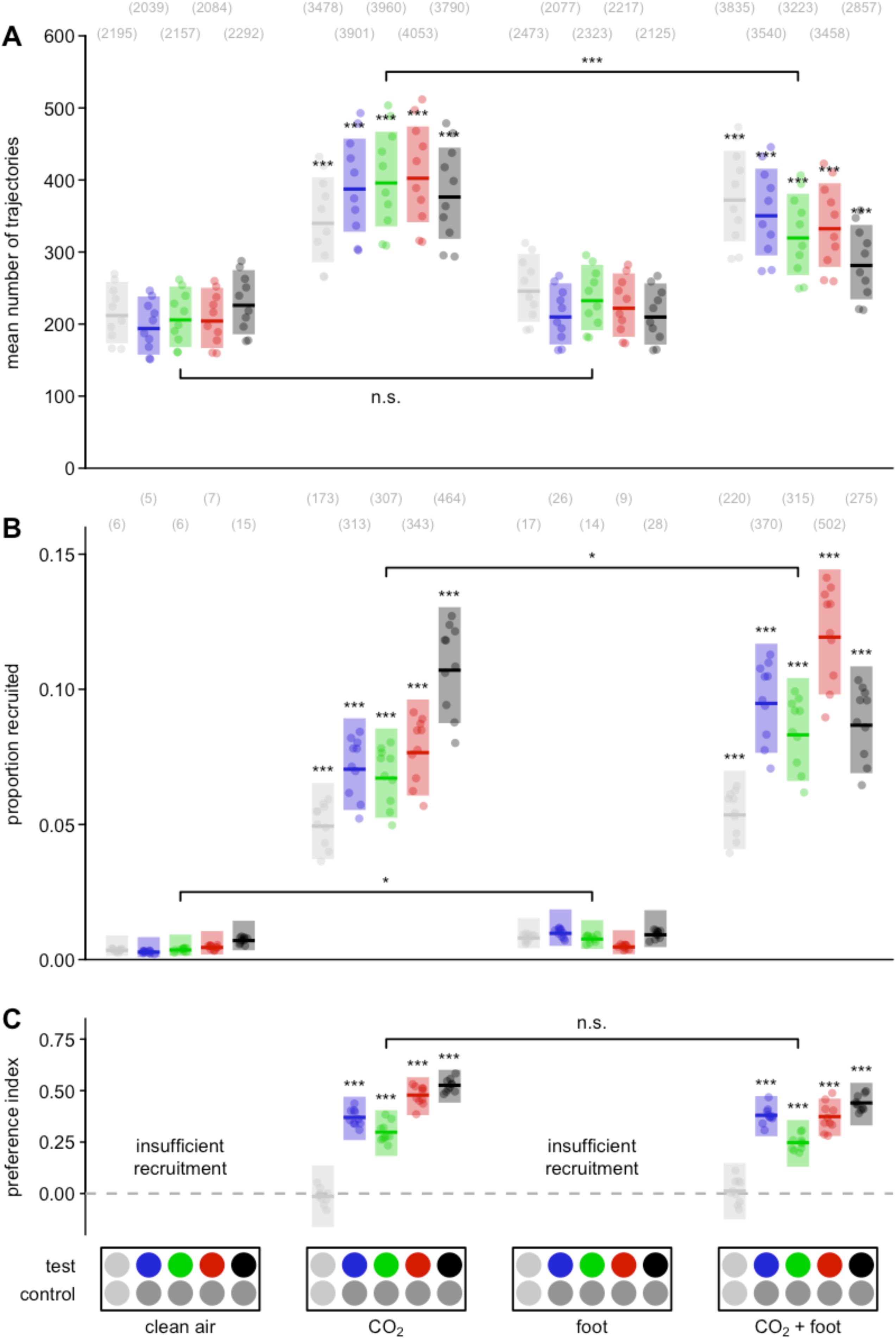
Effect of foot odor on mosquito behavior. **A** The mean number of mosquito trajectories recorded, which measured mosquito activation, during each stimulus period where the plume consisted of clean air alone, 10% CO2, 10% of the air passing through an odor jar containing a source of human foot odor, and the combination of CO2 and foot odor. Asterisks above the boxes here indicate a statistical difference in the number of trajectories as compared with the stimulus period with paired neutral gray stimuli and clean air alone (leftmost box). When CO2 was present, we found foot odor caused a significant decrease in the number of trajectories (significance bracket, *a priori* contrast, z = 3.34, *P* = 0.0008). Bracketed numbers above each boxplot indicate the total number of trajectories over 10 bioassay runs. **B** The proportion of trajectories recruited to either the test or control visual stimuli under the same odor conditions listed above. Asterisks above the boxes denote a statistically significant difference from the recruitment to paired neutral gray stimuli with clean air alone (leftmost box). We found a small, statistically significant, increase in the recruitment between clean air alone and foot odor (significance bracket, *a priori* contrast, z =-2.42, *P* = 0.0153), and a significant increase in the recruitment between CO2 alone and the combination of and CO2 and foot odor (significance bracket, *a priori* contrast, z =-2.48 *P* = 0.0129). Bracketed numbers between panels indicate the number of recruited trajectories over 10 bioassay runs. **C** The preference index of mosquitos in the wind tunnel responding to visual stimuli of various colors under the same odor conditions listed above. Significance stars above the boxes here indicate a difference from a preference index of 0.00. We found no evidence that foot odor in the plume influenced the visual preference of responding mosquitos, at least among this limited set of visual stimuli (significance bracket, likelihood-ratio test, χ^2^ = 4.82 df = 5 *P* = 0.44). Test stimuli from left to right: neutral gray (light gray circles) at an intensity matching the fabric background, blue (450 nm), green (527 nm), red (621 nm) LEDs at matching isoquantal intensities, and unilluminated black tulle targets (black circles). Control stimuli: neutral or mid gray. Boxplots are the mean (line) with 95% confidence interval (shaded area), with points representing model predictions for each of replicate bioassay run. n.s. > 0.05, \**P* < 0.05, \*\**P* < 0.01, \*\*\**P* < 0.001

**Figure S7.**
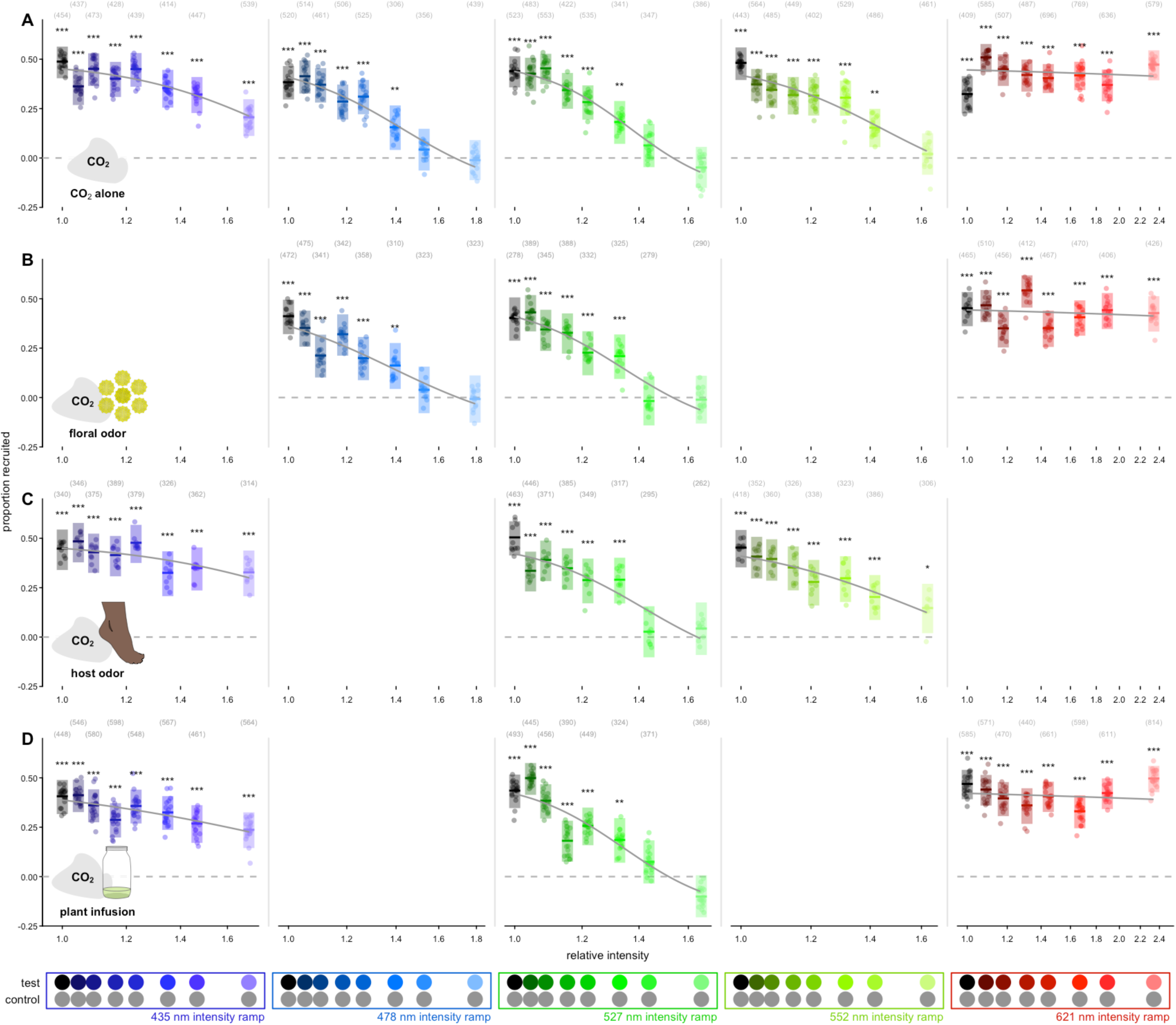
**Effect of odor and stimulus intensity on mosquito visual preference**. Proportion of trajectories recruited to either the test or control visual stimulus in the presence of CO2 paired with (**A**) no odor, (**B**) tansy (*T. vulgare*) floral odor, (**C**) human foot odor, and (**D**) the odor of an alfalfa infusion. We investigated the effect of intensity at a selection of wavelengths covering the visible range and focusing on spectral ranges where we observed odor shifts in spectral preferences in the spectral sweep experiments. The intensities on the x-axes are measured relative to the unilluminated black tulle targets, which were common among all of the intensity ramps, and non-zero due to the ambient illumination. Test stimuli: 435 nm, 470 nm, 527 nm, 552 nm, and 621 nm LED intensity ramps ranging in intensity from 0.0 to 3.0 times the isoquantal intensity used in the spectral sweep experiments. Control stimuli: mid gray. Boxplots are the mean (line) with 95% confidence interval (shaded area), with points representing model predictions for each of replicate bioassay run. Bracketed numbers above each bar indicate the number of recruited trajectories over 20, 16, 10 and 20 replicate bioassay runs respectively. Gray lines show a sigmoid fitted to the preference data. Asterisks above the boxes denote a statistically significant difference from a preference index of 0.00. \**P* < 0.05, \*\**P* < 0.01, \*\*\**P* < 0.001

## Notes

https://doi.org/10.17632/fdr7znz5dh.1

